# Re-entry into mitosis and regeneration of intestinal stem cells through enteroblast dedifferentiation in *Drosophila* midguts

**DOI:** 10.1101/2021.11.22.469515

**Authors:** Aiguo Tian, Virginia Morejon, Sarah Kohoutek, Yi-Chun Huang, Wu-Min Deng, Jin Jiang

**Affiliations:** Department of Biochemistry and Molecular Biology, Tulane University School of Medicine, Louisiana Cancer Research Center, New Orleans, LA 70112, USA; Department of Molecular Biology, University of Texas Southwestern Medical Center, Dallas, TX

**Keywords:** Dedifferentiation, Stem cells, Enteroblasts, Epithermal growth factor receptor signaling, *Drosophila* midgut

## Abstract

Many adult tissues and organs including the intestine rely on resident stem cells to maintain homeostasis. In mammalian intestines, upon ablation of resident stem cells, the progenies of intestinal stem cells (ISCs) such as secretory cells and tuft cells can dedifferentiate to generate ISCs to drive epithelial regeneration, but whether and how the ISC progenies dedifferentiate to generate ISCs under physiological conditions remains unknown. Here we show that infection of pathogenic bacteria induces enteroblasts (EBs) as one type of ISC progenies to re-enter the mitotic cycle in the *Drosophila* intestine. The re-entry into mitosis is dependent on epithermal growth factor receptor (EGFR)-Ras signaling and ectopic activation of EGFR-Ras signaling in EBs is sufficient to drive EBs cell-autonomously to re-enter into mitosis. In addition, we examined whether EBs gain ISC identity as a prerequisite to divide, but the immunostaining with stem cell marker Delta shows that these dividing EBs do not gain ISC identity. After employing lineage tracing experiments, we further demonstrate that EBs dedifferentiate to generate functional ISCs after symmetric divisions of EBs. Together, our study in *Drosophila* intestines uncovers a new role of EGFR-Ras signaling in regulating re-entry into mitosis and dedifferentiation during regeneration and reveals a novel mechanism by which ISC progenies undergo dedifferentiation through a mitotic division, which has important implication to mammalian tissue homeostasis and tumorigenesis.

## Introduction

The *Drosophila* midgut is the functional equivalent of mammalian small intestine where food is digested and nutrients are absorbed, and they protect the internal gut milieu from external environment (Jiang and Edgar, 2012; Li and Jasper, 2016; Sansonetti, 2004; Zwick et al., 2019). During normal homeostasis, both the *Drosophila* midgut and the mammalian small intestine undergo cellular turnover to maintain tissue integrity and function. In response to tissue damage such as chemical feeding or bacterial infection, they can mount regenerative programs to accelerate stem cell division and differentiation to effectively replenish damaged mature cells (Amcheslavsky et al., 2009; Biteau et al., 2011; Jiang and Edgar, 2012; Jiang et al., 2016). ISCs in both mammals and *Drosophila* divide continuously to give rise to stem cells and progenitor cells, which differentiate into enterocytes (ECs) or enteroendocrine (EE) cells or into additional mature cells in mammals (such as Paneth cells, tuft cells and goblet cells) (Barker et al., 2007; Beumer and Clevers, 2016; Jiang and Edgar, 2012). These progenitors from ISCs in mammals enter the trans-amplifying compartment to rapidly divide before terminal differentiation, but the progenitors in *Drosophila* such as EBs which start endoreplication to become ECs cannot divide anymore (Beumer and Clevers, 2016; Micchelli and Perrimon, 2006; Ohlstein and Spradling, 2006). During *Drosophila* intestinal regeneration, EGFR-Ras signaling pathway and other signaling pathways, such as Notch, Wnt, Hh and BMP pathways are implicated in the regulation of ISC self-renewal, proliferation and/or differentiation (Beehler-Evans and Micchelli, 2015; Biteau and Jasper, 2014; Chen et al., 2018; Jiang et al., 2011; Lin et al., 2008; Micchelli and Perrimon, 2006; Ohlstein and Spradling, 2006; Ohlstein and Spradling, 2007; Tian and Jiang, 2014; Tian et al., 2015; Tian et al., 2017; Zeng and Hou, 2015). As a result of activation of EGFR-Ras signaling in progenitors cells including ISCs and EBs, ISCs enter the mitosis quickly to speed up proliferation and regeneration (Biteau and Jasper, 2011; Buchon et al., 2010; Jiang et al., 2011). However, whether EBs as the ISC progenies could re-enter mitosis through EGFR-Ras signaling remains largely unexplored.

In the mammalian intestine, the progenitors, such as secretory progenitors and enteroblast progenitors, and mature cells, such as secretory cells, tuft cells and Paneth cells, could dedifferentiate to produce ISCs after ablation of ISCs (Jones et al., 2019; Murata et al., 2020; Schmitt et al., 2018; Tetteh et al., 2016; van Es et al., 2012; Yu et al., 2018). Several signaling pathways including Notch have been shown to be involved in the dedifferentiation. However, the process of dedifferentiation and the underlying molecular and cellular mechanisms remain poorly understood. Until now, only resident intestinal stem cells (ISCs) and regenerated ISCs from differentiating ECs through amitosis have been reported in *Drosophila* midguts (Lucchetta and Ohlstein, 2017; Micchelli and Perrimon, 2006; Ohlstein and Spradling, 2006), but whether and how EBs dedifferentiate to replenish the ISC pool remains unknown.

In this study, we explore whether and how EBs as the immediate progenies of ISCs re-enter the mitotic cycle during intestinal regeneration in response to bacterial infection in *Drosophila*. Our study provides evidence that EBs can undergo mitosis and dedifferentiation under physiological challenges and uncover a novel role of EGFR-Ras signaling in EBs in the cell-autonomous regulation of re-entry into mitosis and dedifferentiation.

## Results

### Bacterial infection induces the re-entry into mitosis in EBs

To determine whether EBs as one type of the progenies of ISCs in *Drosophila* midguts can re-enter the mitotic cycle upon microbial infection, adult female flies were fed with pathogenic bacteria (*Pseudomonas entomophila*, *P. e*) for 36 hours (h), and mitosis in midguts was examined with the mitotic marker (phospho-Ser10-Histone H3, PH3). The EBs, which normally enter endoreplication and differentiate into ECs, are marked by expression of *UAS-GFP* driven by *Su(H)-Gal4* (a EB specific Gal4 (Zeng et al., 2010) and *Su(H)–Gal4*>*UAS–GFP* is referred to as *Su(H)*>*GFP*) (Figure 1A). Flies fed with sucrose (suc) were used as the control. We found that the midguts from female flies with *P. e* infection or the control showed PH3 staining in ISCs (PH3^+^GFP^−^, Figure 1B-C’, D-E’, white arrowheads), and the number of PH3^+^ ISCs was significantly increased in *P .e* infected midguts compared to those of the control (Figure 1J: 64.3 PH3/gut in *P. e* vs. 5.4 PH3/gut in the control; n=12), as previously reported (Jiang et al., 2011). Interestingly, PH3 staining was also found in some EBs in *P. e* infected midguts (PH3^+^GFP^+^, Figure 1D, F-F’, red arrows), but not in the control midguts (Figure 1B) (Figure 1K: 8.13 PH3/gut in *P. e* vs. 0 PH3/gut in the control, n=12), indicating that *P. e* infection can induce re-entry into mitosis in EBs. When feeding adult female flies with another pathogenic bacteria (*Erwinia carotovora carotovora* strain *15*, *Ecc15*), both EB mitosis (PH3^+^GFP^+^ cells, Figure 1G, I-I’, red arrows, K; 5.5 PH3/gut, n=12) and significant increase of ISC mitosis (PH3^+^GFP^−^, Figure 1G-H’, white arrowhead, J; 60.75 PH3/gut, n=12) were observed in their midguts. The re-entry into mitosis in EBs induced by either *P. e* or *Ecc15* infection suggested the general effect of pathogenic bacteria on inducing ISC mitosis and EB re-entry into mitosis in the intestine. To rule out the leaky expression of *Su(H)*>*GFP* in ISCs, we fed flies with dextran sodium sulfate (Dss) which damages the gut basement membrane to induce ISC proliferation (Amcheslavsky et al., 2009; Ren et al., 2013; Tian et al., 2015) and found increased number of ISCs (GFP^−^) with PH3, but no EBs (GFP^+^) with PH3 (Figure 1—figure supplement 1).

**Figure 1.**
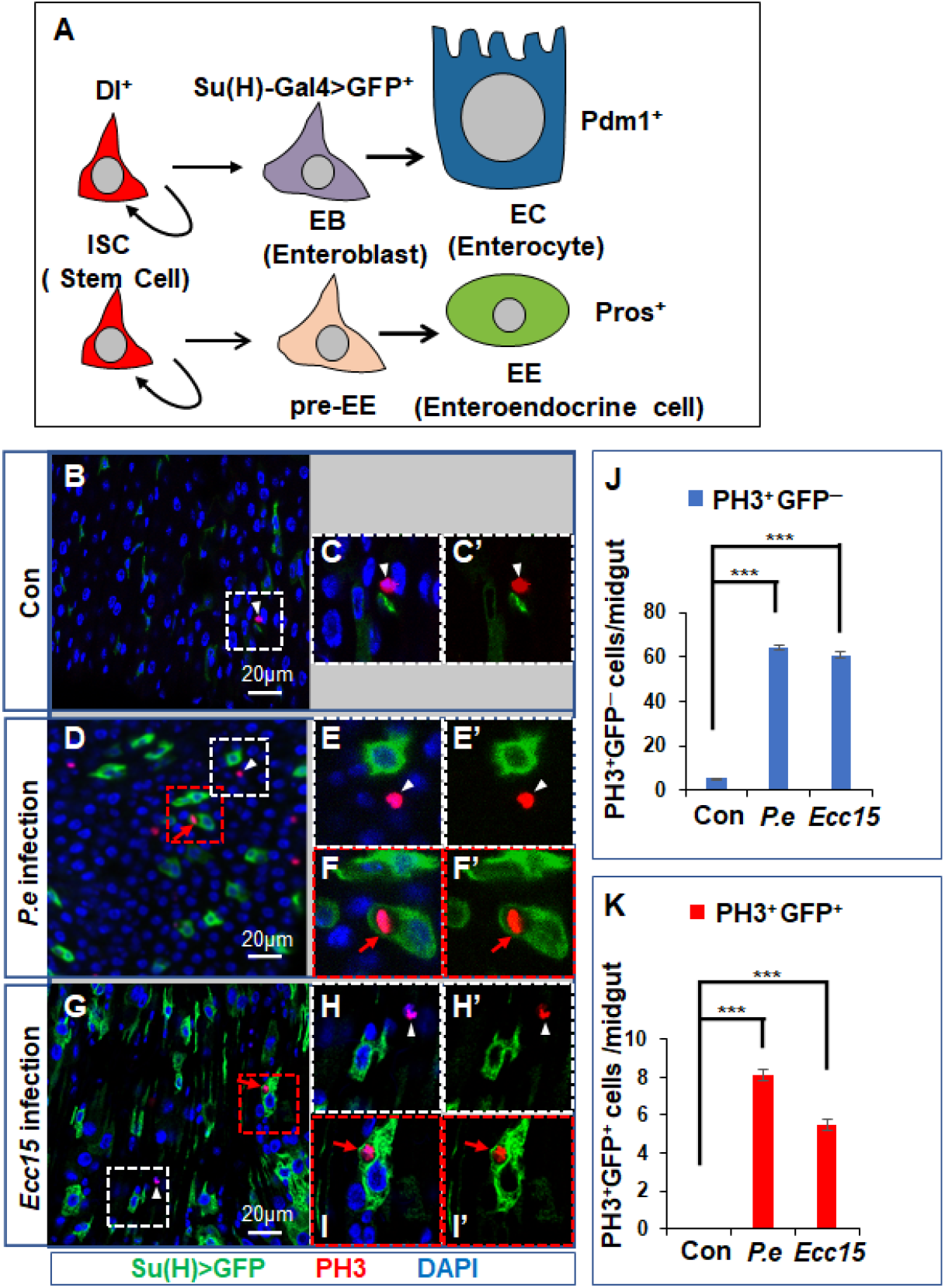
Bacterial infection can induce re-entry into mitosis in EBs. (A) ISC lineages in *Drosophila* adult midguts. Dl marks ISCs. *Su(H)-LacZ* or *Su(H)-Gal4>UAS-GFP* marks EBs. Pdm1 and Prospero (Pros) are markers for EC and EE, respectively. ISC, EB and pre-enteroendocrine (pre-EE) cells are collectively called progenitors cells. (B-I’) The *Drosophila* midguts containing *Su(H)-Gal4*>*UAS-GFP* with sucrose (suc, control) (B-C’), *P.e* feeding for 36 h (D-F’) and *Ecc15* feeding for 36 h (G-I’) were immunostained for GFP (green), PH3 (red) and DAPI (blue). (C-C’, E-E’ and H-H’) Magnification of selecting areas containing PH3^+^GFP^−^ cells from B, D and G. (F-F’ and I-I’) Magnification of selecting areas containing PH3^+^GFP^+^ cells from D and G. White arrowheads and red arrows indicate PH3 in GFP^−^ and GFP^+^ cells, respectively. (J and K) Quantification of PH3^+^GFP^−^ ISCs (J) and PH3^+^GFP^+^ EBs (K) with given treatments. n = 12 guts for each treatment. Three independent experiments were performed, and error bars are ± SEM. ***, P < 0.001 (Student’s t-test).

### EGFR-Ras signaling in EBs is up-regulated upon bacterial infection

Bacterial infection activates signaling pathways such as EGFR-Ras pathway through up-regulation of ligands (Jiang et al., 2011) to induce ISC proliferation. To assess the expression level of ligands of the EGFR-Ras pathway, we performed qRT-PCR analysis and confirmed that *P. e* infection upregulated expression of the ligand *Vn* and *Krn* (Figure 2A). To determine whether EGFR-Ras signaling was up-regulated in EBs upon *P. e* infection, we examined the activity of mitogen-activated protein kinase (MAPK), which marks the activation of EGFR-Ras signaling, by using an antibody against the diphosphorylated and active form of MAPK (dpERK) (Gabay et al., 1997). We found that the dpErk level was high in GFP^−^ ISCs (Figure 2B-B”, arrowheads) but low in GFP^+^ EBs (Figure 2B-B”, arrows) in the control midgut, as previously reported (Jiang et al., 2011). After *P. e* infection, the level of dpERK was greatly increased in both GFP^+^ EBs (Figure 2C-C”’, arrows) and GFP^−^ ISCs (Figure 2C-C”’, arrowheads), suggesting that bacterial infection activates EGFR-Ras signaling strongly in EBs, as well as in ISCs.

**Figure 2.**
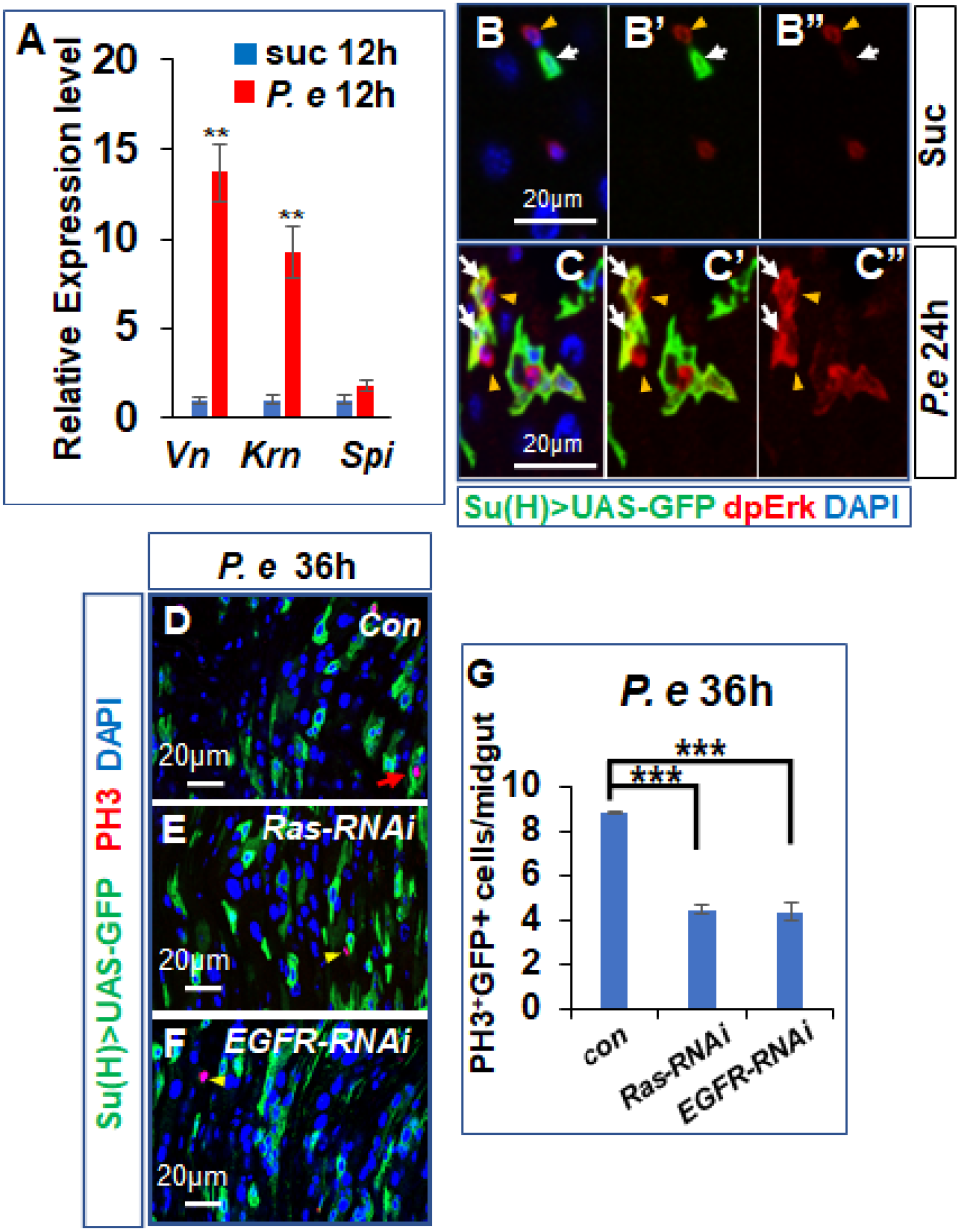
EGFR-Ras signaling is required for EB mitosis up bacterial infection. (A) *P. e* infection up-regulates the expression of EGFR ligand *Vn* and *Krn*. Three independent experiments were performed, and error bars are ± SEM. **, P < 0.01. (B-B”) In the control, the dpErk is expressed in ISCs (arrowheads), and weakly expressed in EBs (arrows). (C-C”) The level of dpErk in *P. e* infected flies is greatly up-regulated in both EBs (arrows) and ISCs (arrowheads). (D-F) The *P. e* infected midguts expressing *Su(H)^ts^*>*GFP* with control (D), *UAS-Ras-RNAi* (E), *UAS-EGFR-RNAi* (F). (G) Quantification of PH3^+^EBs (GFP^+^) with the given genotypes for *P. e* infection. n = 11 guts for each genotype. Three independent experiments were performed, and error bars are ± SEM. ***, P < 0.001 (Student’s t-test).

### EGFR-Ras signaling in EBs is required for bacterial infection induced EB mitosis

The up-regulation of EGFR-Ras signaling in EBs by *P. e* infection led us to hypothesize that activation of EGFR-Ras signaling in EBs may induce EB mitosis. To test this, we activated EGFR-Ras signaling in EBs and examined the expression of PH3. The inducible Gal4/Gal80^ts^ system with EB specific Gal4 (*Su(H)-Gal4 Tub-Gal80^ts^*; referred to as *Su(H)^ts^*) was used to overexpress the active form of *Ras* (*UAS-Ras^V12^*) (*Karim and Rubin, 1998*) in EBs. For these experiments with Gal80^ts^, female flies expressing *Su(H)^ts^*>*UAS-GFP* with or without *UAS–gene* were raised to adults at 18°C (Gal4 is ‘off’) and then shifted to 29°C to degrade Gal80^ts^ (Gal4 is ‘on’) for indicated days so that *Su(H)-Gal4* can drive expression of *UAS-gene*. Our assay with expression of *Ras^V12^* in EBs found PH3 in GFP^+^ EBs, indicating that activation of *Ras* in EBs induced cell-autonomous EB mitosis (GFP^+^ PH3^+^, Figure 3C, E, E’, L, red arrows; 12.7 PH3/gut in *Ras^V12^* vs. 0 PH3/gut in the control, n=13). In addition, we found that activation of *Ras* in EBs non-cell-autonomously promote ISC proliferation (PH3^+^ GFP^−^, Figure 3C-D’, arrowheads, Figure 3M; 27.8 PH3/gut in *Ras^V12^* vs. 5.4 PH3/gut in the control, n=13), indicated by increased number of PH3 in GFP^−^ cells. To further confirm that EGFR-Ras signaling can induce EB mitosis, we used the same system to overexpress the active form of *EGFR* (*EGFR^A887T^* or λ*Top*) (Lesokhin et al., 1999; Queenan et al., 1997) in EBs, and found that overexpression of either *EGFR^A887T^* or λ*Top* at 29°C for 4d or 5d also induced EB mitosis (PH3^+^GFP^+^; Figure 3F, H, H’, I, K, K’, red arrows, Figure 3L; 9.9 PH3/gut in *EGFR^A887T^* vs. 6.6 PH3/gut in λ*Top* vs. 0 PH3/gut in the control, n=13) and non-cell-autonomously increased ISC proliferation (GFP^-^ PH3^+^, Figure 3F-G’, I-J’, arrowheads, Figure 3M). To exclude the leaky expression of *Su(H)-Gal4* in ISCs, we used *Su(H)^ts^*>*GFP* to drive expression of an active form of *InR* (*UAS-InR^ACT^*) in EBs and found increased number of PH3 in GFP^−^ ISCs, but no PH3 in GFP^+^ EBs (Figure 3—figure supplement 1).

**Figure 3.**
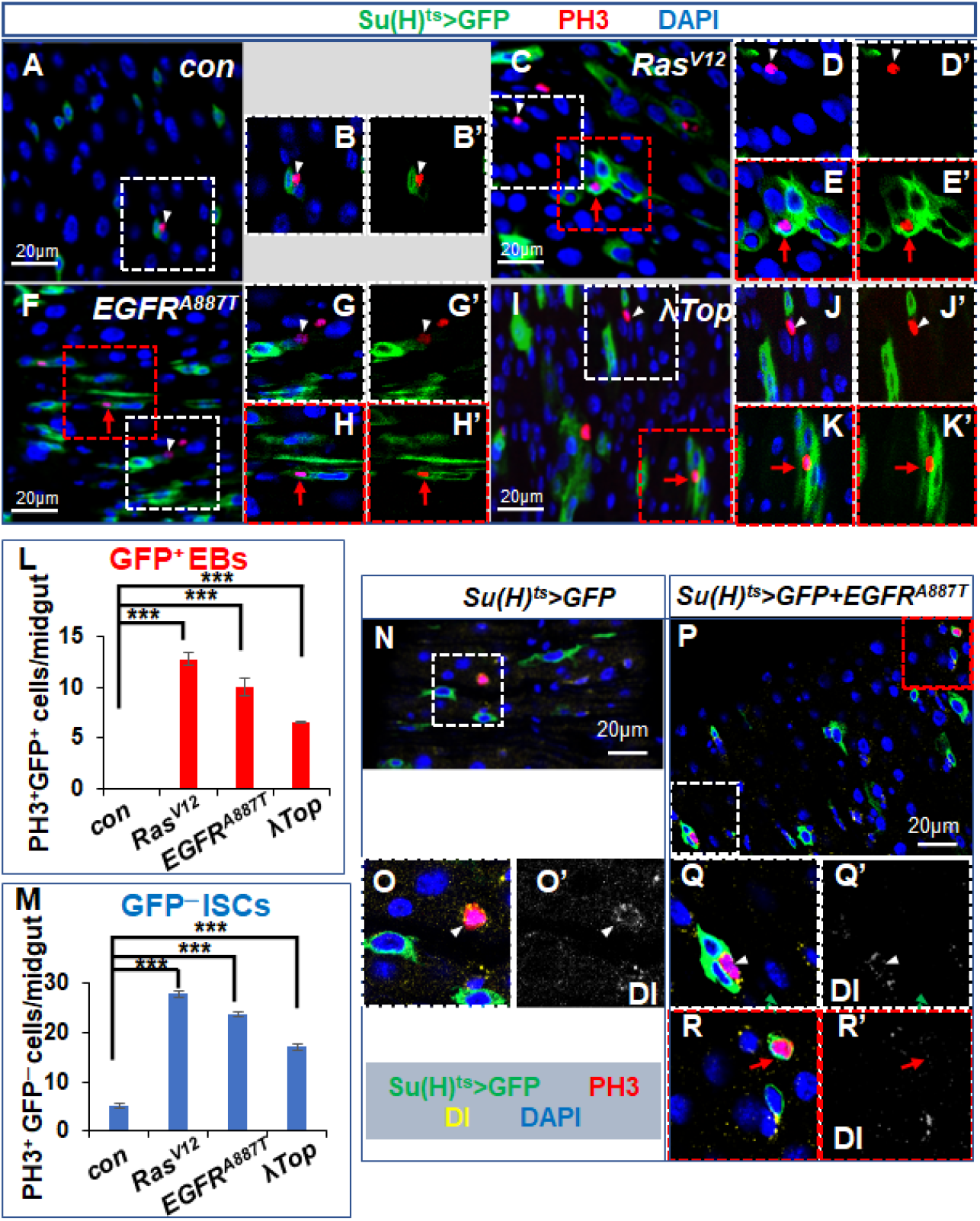
Activation of EGFR-Ras signaling in EBs induces cell-autonomous EB mitosis and non-cell-autonomously promotes ISC proliferation. (A-K’) The *Drosophila* midguts expressing *Su(H)^ts^*>*GFP* with control (A-B’), *UAS-Ras^V12^* for 6 days (C-E’), *EGFR^A887T^* for 4 days (F-H’) and *UAS*-λ*Top* for 5 days (I-K’) were immunostained for GFP (green), PH3 (red) and DAPI (blue). (B-B’, D-D’, G-G’ and J-J’) Magnification of selecting areas containing PH3^+^GFP^−^ cells from A, C, F and I. (E-E’, H-H’ and K-K’) Magnification of selecting areas containing PH3^+^GFP^+^ cells from C, F and I. White arrowheads and red arrows indicate PH3 in GFP^−^ and GFP^+^ cells, respectively. (L, M) Quantification of PH3^+^ ISCs (GFP^−^, M) and PH3^+^ EBs (GFP^+^, L) in midguts with indicated genotypes. n = 13 guts for each genotype. Three independent experiments were performed, and error bars ± SEM. ***, P < 0.001 (Student’s t-test). (N-R’) PH3 and Dl staining in the control (N-O’) or *EGFR^A887T^* overexpressed midguts (P-R’). The white arrowheads indicate that GFP^−^ ISCs with PH3 have Dl staining (O-O’, Q-Q’); the red arrows indicate that EBs with PH3 and GFP (GFP^+^PH3^+^) do not have Dl expression (R-R’).

To confirm that these GFP^+^ EBs did not gain ISC identity as a prerequisite to divide, we performed the immunostaining with ISC marker (Delta, Dl) and found that these PH3^+^ ISCs without GFP did have Dl expression (Figure 3N-O’, P-Q’), but these EBs marked by Su(H)>GFP with PH3 did not have Dl expression (Figure 3P, R-R’, 100%, n=42), suggesting that these dividing EBs do not gain ISC identity. In summary, activation of EGFR-Ras signaling in EBs induces cell-autonomous re-entry into mitosis in EBs and non-cell-autonomously promotes ISC proliferation.

To determine whether EGFR-Ras signaling in EBs is necessary for *P. e* infection induced re-entry into mitosis in EBs, we inactivated the EGFR-Ras signaling in EBs by knocking down either *EGFR* or *Ras* with *UAS-EGFR-RNAi* (or *UAS-Ras-RNAi*) and *Su(H)^ts^* (referred to as *Su(H)^ts^*>*EGFR-RNAi* (or *Ras-RNAi*)) and analyzed the consequence of EB mitosis in response to *P. e* infection. The knockdown of either *EGFR* or *Ras* decreased EB mitosis, as indicated by the reduced number of PH3^+^ cells in EBs compared with the control (Figure 2D-G: 8.9 PH3/gut in control vs. 4.5 PH3/gut in *EGFR-RNAi* vs. 4.4 PH3/gut in *Ras-RNAi;* n=11), indicating that EGFR-Ras signaling in EBs is required for EB mitosis upon bacterial infection.

### New Dl^+^ ISCs are generated from EBs upon *P. e* infection

Upon bacterial infection, EBs re-enter the mitotic cycle to divide and produce new cells, thus we asked whether these new cells could become ISCs. To test this, we used the lineage tracing system (referred to as Flip-out system, Figure 4A) (Germani et al., 2018; Golic and Lindquist, 1989) to trace the progenies of EBs. In this lineage tracing system, the expression of flippase (Flp) driven by *Su(H)-Gal4* induces the flipping out of the FRTs in the *actP (FRT Stop FRT) LacZ* cassette (referred to as *actP*>*stop*>*lacZ*), thus *actP-LacZ* was produced and expressed in EBs and their progenies. For the lineage tracing experiment with feeding of sucrose, the immunostaining with antibodies against GFP, β-gal and Delta (Dl, the ISC marker) showed that *LacZ* expression was found in GFP^+^ EBs (*GFP*^+^*LacZ*^+^, Figure 4B-B’, arrows), but not in Dl^+^ GFP^−^ ISCs (Fig.4B-B”’, blue arrowheads). In contrast, after feeding with *P. e*, *LacZ*^+^ was found to be expressed in cells with GFP (Figure 4C-C”’, arrows) or without GFP (Figure 4C-C”’, red arrowheads). These *LacZ*^+^ cells without GFP indicated that they were progenies of EBs but were not EBs anymore. In these *LacZ*^+^*GFP*_−_ cells, Dl was found in some of them (Figure 4C-C”’, red and blue arrowheads, Figure 4D), indicating that these ISCs (*LacZ*^+^*GFP*^−^*Dl*^+^) were generated from EBs. The other *GFP*^−^*LacZ*^+^ cells (Figure 4C-C”‘, only red arrowhead) without Dl staining should be differentiating ECs from EB differentiation. In summary, these results indicate that *P. e* infection could induce EB dedifferentiation to generate ISCs.

**Figure 4.**
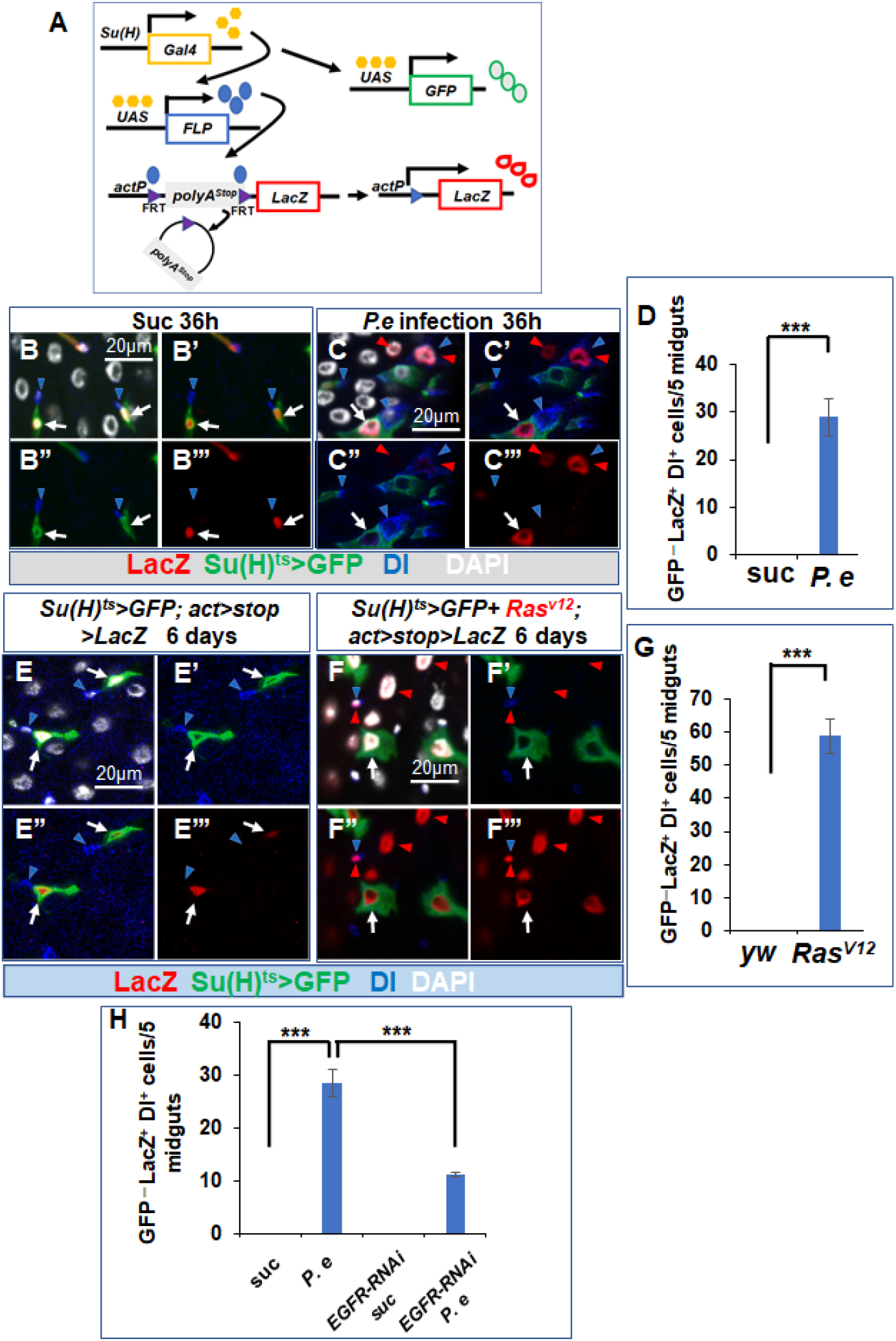
Dl^+^ ISCs are generated from EBs upon *P.e* infection and activation of EGFR-Ras signaling in EBs. (A) The schematic drawing of the Flip-out lineage tracing system. (B-C”’) Flies expressing *UAS-Flp; Su(H)^ts^ UAS-GFP*; *ActP*>*Stop*>*LacZ* were fed with Suc (B-B”’) or *P.e* (C-C”’) for 36h and their midguts were dissected and immunostained with antibodies to GFP, β-gal and Dl. (D) Quantification of *GFP*^−^*LacZ*^+^*Dl*^+^ in the control, *P. e* infected midguts. n=50. Three independent experiments were performed, and error bars are ± SEM. ***, P < 0.001 (Student’s t-test). (E-F”’) The midguts from flies expressing *UAS-Flp*; *Su(H)^ts^ UAS-GFP*; *ActP*>*Stop*>*LacZ* with (F-F”’) or without (E-E”’) *UAS-Ras^V12^* were dissected and immunostained with antibodies to GFP, β-gal and Dl. (G) Quantification of *GFP*^−^*LacZ*^+^*Dl*^+^ with the given genotypes. n=50. Three independent experiments were performed, and error bars are ± SEM. ***, P < 0.001 (Student’s t-test). (H) Quantification of *GFP*^−^*LacZ*^+^*Dl*^+^ in the control with suc or *P. e*, or knockdown of *EGFR* with suc or *P. e* infection. n=50. Three independent experiments were performed, and error bars are ± SEM. ***, P < 0.001 (Student’s t-test).

### Activating EGFR-Ras signaling in EBs triggers EB dedifferentiation to generate Dl^+^ ISCs

The observation that *P.e* infection induces EB dedifferentiation suggests that activation of EGFR-Ras signaling in EBs may induce EB dedifferentiation. Indeed, after we performed the lineage tracing experiments with the same Flip-out system and overexpression of *UAS-Ras^V12^* in EBs for 6d, we found LacZ^+^ EB cells with GFP (*GFP*^+^*LacZ*^+^, Figure 4F-F”’, arrows) and LacZ^+^ cells without GFP (*GFP*^−^*LacZ*^+^, Figure 4F-F”’, red arrowheads). In these *GFP*^−^*LacZ*^+^ cells, some showed Dl expression (*GFP*_−_*LacZ*^+^*Dl*^+^) (Figure 4F-F”’, red and blue arrowheads, and 4G), indicating that activation of EGFR-Ras signaling in EBs can induce EB dedifferentiation to generate ISCs.

To determine whether *P. e* infection induces EB dedifferentiation through EGFR-Ras signaling. We inactivated the EGFR-Ras signaling in EBs by knocking down *EGFR* with the Flp-out system and examined whether the new regenerated ISCs were blocked upon *P.e* infection. We found that the total number of regenerated ISCs from EBs was reduced with EGFR knockdown in response to *P. e* infection (total number of *GFP*^−^*LacZ*^+^*Dl*^+^ cells in five guts, 28.5 in *P. e* infection vs. 11.3 in *EGFR-RNAi* with *P. e* infection, Figure 4H).

Our next question is whether these new Dl^+^ stem cells from EB dedifferentiation are functional. To test this, differentiation of ISCs into EE cells marked by expression of *Prospero* (*Pros*) was used as a readout for functional ISCs. After activating EGFR-Ras signaling in EBs with *Ras^V12^* or *EGFR^A887T^* expression for 6d or *P.e* infection for 36h, we found that some LacZ expressed cells without GFP have gained Pros expression (Figure 5A-D), indicating that these ISCs from EB dedifferentiation are functional so that mature EE cells are generated.

**Figure 5.**
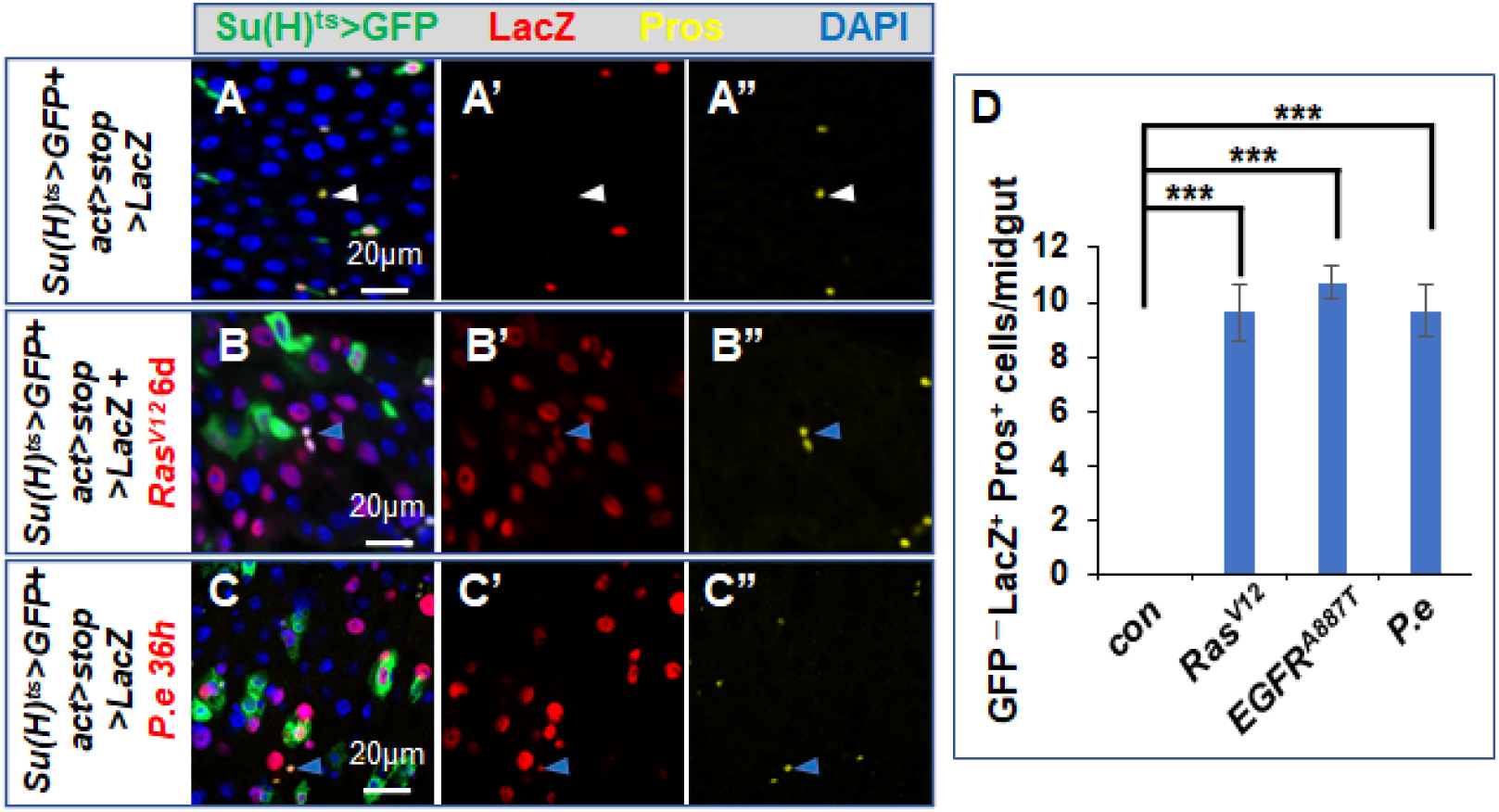
The regenerated ISCs from EBs can generate EE cells. (B-C”) ISCs from EBs with *Ras^V12^* expression or *P. e* infection could differentiate to generate EE cells (*LacZ*^+^*Pros*^+^*GFP*^−^, blue arrowheads), but no EE cells from EBs are found in the control (A-A”, white arrowheads). (D) Quantification of EE cells from progenies of EBs with indicated genotypes or treatments. n = 9 guts for each genotype or treatment. Three independent experiments were performed, and error bars are ± SEM. ***, P < 0.001 (Student’s t-test).

### The symmetric division of EBs produces two ISCs upon activation of EGFR-Ras signaling in EBs

Our results show that EBs can re-enter mitosis and ISCs are generated from EBs, and the next question is whether EBs generate ISCs after mitosis or EBs gain stem cell identity before mitosis. Firstly, we excluded the possibility that these dividing EBs have gained ISC property marked by Dl expression (Figure 3P, R-R’). Secondly, we compared mitosis in EBs to that in ISCs by examining the centrosome marker and different mitotic stages. The results showed that similar mitosis with different stages were found in EBs (Figure 6D-I), as well as in ISCs (Figure 6A-C). At last, we performed the two-color lineage tracing experiment which is based on the mitotic division of cells (Figure 6J, K) to trace the lineage cells from an individual EB division. In this experiment, 3-5-day-old flies expressing *UAS-Flp*; *Su(H)^ts^*; *FRT82BGFP/FRT82BRFP* with or without *UAS-EGFRA^887T^* raised in 18°C was transferred to 29°C for 5 days and these pairs of single-color clones with GFP or RFP in the midguts were analyzed (Figure 6L-N). We found that pairs of single-color clones were produced in flies with activation of *EGFR-Ras* signaling with *UAS-EGFR^A887T^* (Figure 6M-N, Figure 6—figure supplement 1), but not in control flies (Figure 6L-L”). This result indicated that EBs with *EGFR^A887T^* expression underwent mitosis, but EBs in the control did not. Interestingly, after the number of cells in single-color clones was counted, we found that the clones in most of pairs of single-color clones had more than one cell (Figure 6M-M”, N), indicating that EBs underwent symmetric cell division to produce two ISCs. In addition, we found low frequency of clones with asymmetric cell division, as indicated by one clone with one cell and the other clone with multiple cells in the pair of single-color of clones (Figure 6—figure supplement 1B-B”, Figure 6N), or with symmetric non-ISC division, as marked by only one cell in each of the pair of single-color of clones (Figure 6—figure supplement 1A-A”, Fig. 6N).

**Figure 6.**
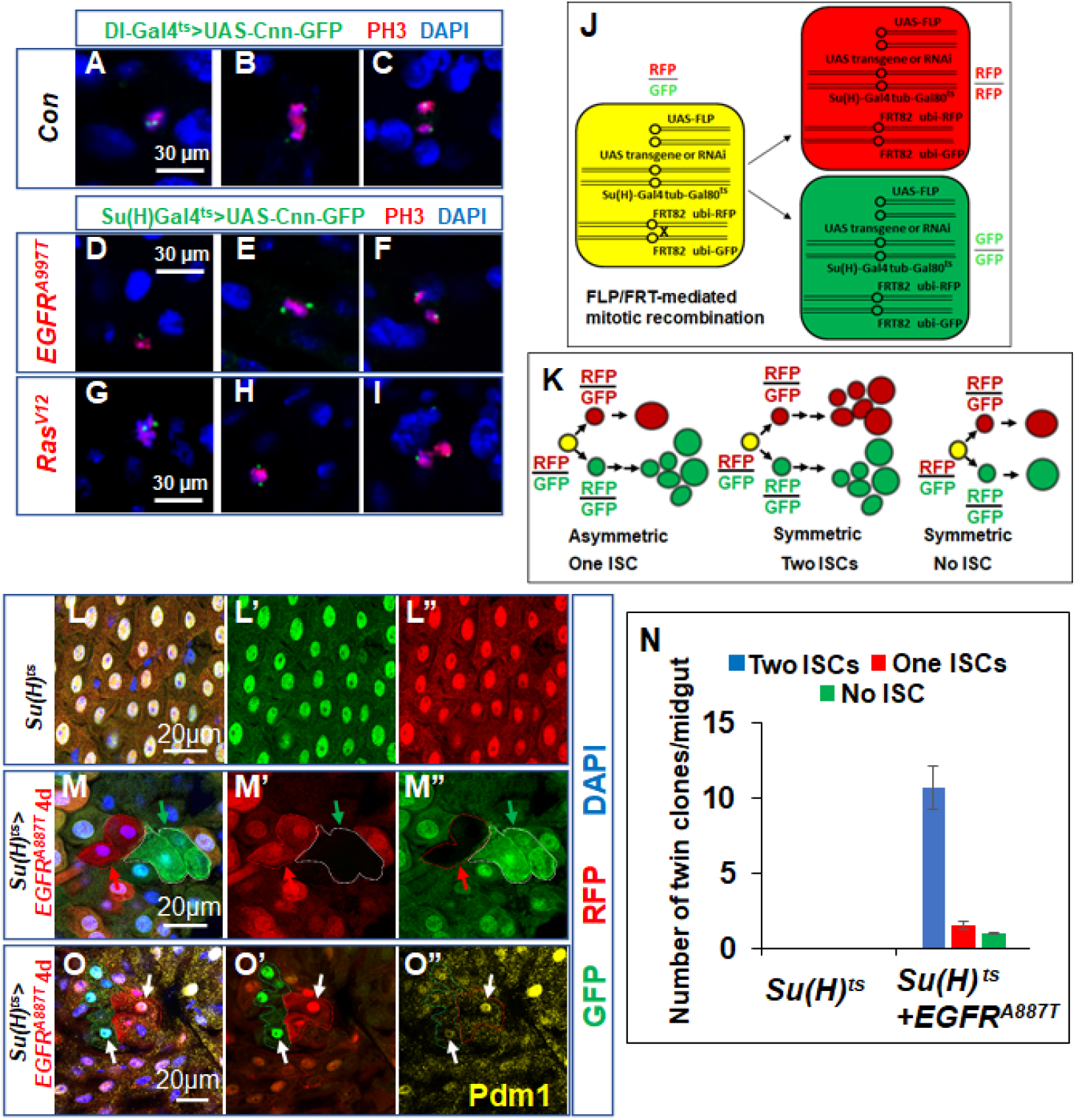
Activating EGFR-Ras signaling in EBs induces symmetric division to generate two functional ISCs. (A-C) The different mitotic stages in ISCs with the centrosome marker (cnn-GFP) which is driven by Dl-Gal4 and PH3. (D-I) The different mitotic stages in EBs with the centrosome marker (cnn-GFP) which is driven by Su(H)-Gal4 and PH3 when *EGFR^A887T^* (D-F) or *Ras^V12^* (G-I) is overexpressed in EBs. (J) Schematic drawing of the cell division that produces differentially labeled twin-spot cells (RFP^+^ GFP^−^ and RFP^−^GFP^+^) through FRT-mediated mitotic recombination. UAS-Flp and transgenic overexpression is driven by *Su(H)^ts^*. (K) Schematic drawings of differentially labeled twin-spot clones generated by FLP/FRT-mediated mitotic recombination of dividing cells in EBs. The representative twin clones from midguts when *EGFRA^887T^* is overexpressed in EBs (M-M”, red and green arrows), but no twin clone is found in the control (L-L”). (N) Quantification of different types of twin clones in the control and *EGFR^A887T^* overexpressed midguts. n = 10 guts for each genotype. Three independent experiments were performed. (O-O”) ECs which were marked by Pdm1 expression were found in pairs of single-color clones (white arrows).

The Flp-out lineage tracing system cannot distinguish ECs from dedifferentiated ISCs or resident ISCs, thus we performed the immunostaining with anti-Pdm1 antibody which mark ECs in the two-color lineage tracing experiments. We found that these dedifferentiated ISCs can differentiate to generate mature ECs with Pdm1 staining (Figure 6O-O”, 100% in single-color clones with multiple cells, n=31), which further indicates that these dedifferentiated ISCs are multipotent to self-renew and generate mature cells.

## Discussion

Previous studies found that only resident ISCs in *Drosophila* midguts localized at the basal side of the gut epithelium undergo asymmetric cell division to produce renewed ISCs and EBs (Micchelli and Perrimon, 2006; Ohlstein and Spradling, 2006). Our studies showed that EBs can dedifferentiate to generate ISCs (Figure 7). There are two possibilities about the process of dedifferentiation with mitosis. One could be that EBs directly revert to ISCs, like induced pluripotent stem cells (Takahashi and Yamanaka, 2006), and then start mitosis. The other one could be that EBs re-enter mitosis, and then dedifferentiate to produce ISCs. Our results from immunostaining with Dl support the latter, and the two-color lineage tracing experiments further demonstrate that two regenerated ISCs are generated from one division of EBs. This process of dedifferentiation is distinct from amitosis, another dedifferentiation process in the intestine, which was found in re-fed condition in starved *Drosophila* midguts to produce ISCs through reduction of ploidy (Lucchetta and Ohlstein, 2017). Therefore, activating EGFR-Ras signaling in EBs firstly enforces these cells to re-enter the mitotic cycle, and then both EGFR-Ras signaling and mitosis may play a role in the fate determination towards ISCs. As dedifferentiation has emerged as a conserved mechanism underlying the replenishment of stem cell pool, dedifferentiation through mitosis could be a conserved mechanism from *Drosophila* to mammals.

**Figure 7.**
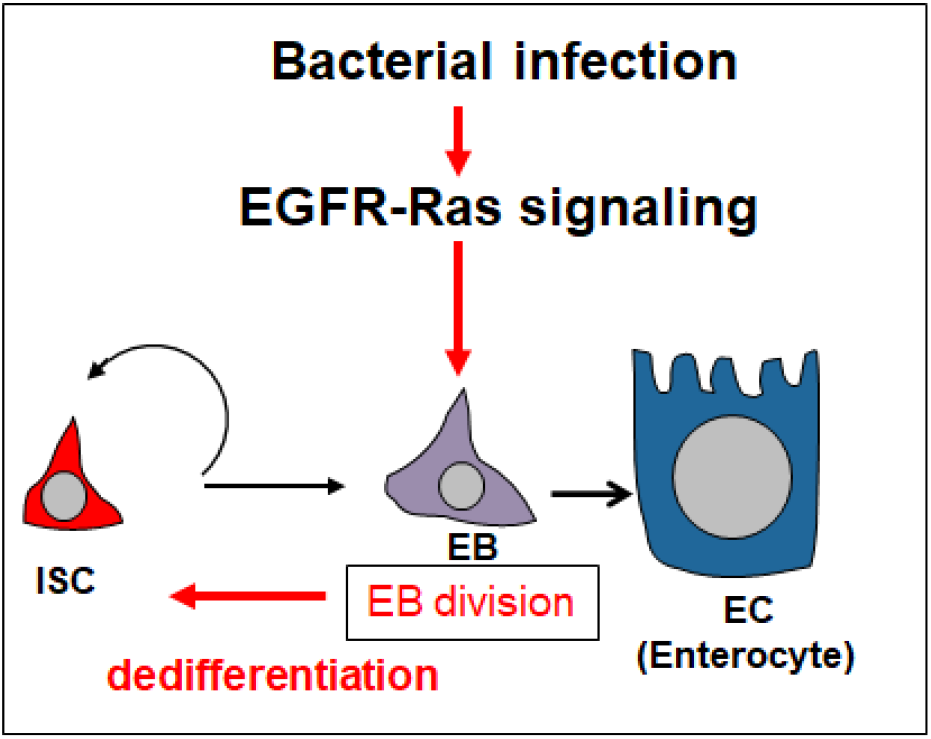
A Model for bacterial infection and activation of EGFR-Ras signaling to induce EB mitosis and dedifferentiation. Bacterial infection induces EGFR ligand expression and activate EGFR-Ras signaling in EBs which is required for EB mitosis. Activating EGFR-Ras signaling in EBs can induce EB dedifferentiation to generate functional ISCs through symmetric division.

Infection of two Gram-negative bacteria induces the same EB mitosis phenotype, indicating that the immunity signaling pathways might be involved in EB mitosis and dedifferentiation. The previous studies showed that bacterial infection could induce the IMD pathway in the *Drosophila* intestine (Buchon et al., 2009; Zhai et al., 2018), but another study reported that Ras/MAPK pathway suppresses IMD in the intestine and fat body (Ragab et al., 2011). Therefore, it is unlikely that EGFR-Ras signaling regulates EB dedifferentiation through IMD pathway. Both bacterial infection and activation of EGFR-Ras signaling can activate JAK-STAT pathway in ISCs through up-regulation of ligands to stimulate ISC proliferation (Jiang et al., 2011; Jiang et al., 2009; Zhai et al., 2018). In addition, JAK-STAT signaling was found to be sufficient to induce dedifferentiation of spermatogonia into germline stem cells in the *Drosophila* testis (Brawley and Matunis, 2004). Thus, EGFR-Ras signaling may regulate EB dedifferentiation through JAK-STAT pathway. Further studies will aid our understanding of the mechanisms underlying intestinal regeneration.

Dysplasia in the mammalian gastrointestinal tract, which is considered as a carcinoma precursor (Li and Jasper, 2016; Pulusu and Lawrance, 2017) and characterized by atypical cellular features, aberrant cell proliferation and differentiation as well as disorganized architecture, can be mimicked in the *Drosophila* intestine (Apidianakis et al., 2009; Biteau et al., 2008; Jiang et al., 2009; Micchelli and Perrimon, 2006; Ohlstein and Spradling, 2006). Upon bacterial infection, the *Drosophila* intestine undergoes rapid regeneration to replace dying cells and this rapid regeneration can be subverted towards dysplasia (Apidianakis et al., 2009; Jiang et al., 2009). During regeneration, ISCs with increased proliferation and increased pool size produce more differentiated cells to replace the lost or damaged cells (Apidianakis et al., 2009; Jiang et al., 2009). Contrary to previous reports, our results indicate that both ISCs and EBs can enter the mitotic cycle, which could make regeneration more efficient. We also find that EBs can generate functional ISCs in response to injury or oncogenic pathway activation, which may contribute to the increased pool of ISCs. Previous studies showed that *Kras* with *Apc* and activation of NF-κB (Schwitalla et al., 2013) acts as a driving force to promote tumorigenesis by dedifferentiation in mammalian colorectal cancer model (Janssen et al., 2006), so, it would be interesting to determine whether the dedifferentiation is mediated by mitosis in mammalian intestines.

## Material and Methods

### *Drosophila* Genetics and Transgenes

Transgenic lines included *UAS-EGFR-RNAi* (VDRC43267), *UAS-EGFRA^887T^* (BL#9534), *UAS-Ras^V12^* (Jiang et al., 2011), *UAS-Ras-RNAi* (BL#34619), *UAS-InR^ACT^* (BL#8440), *UAS-Flp* (BL#55808) *FRT82BUbi-GFP/CyO*, *FRT82BUbi-RFP/CyO*, *Su(H)-Gal4*, *UAS-FLP*; *Su(H)-Gal4 tub-Gal80^ts^ UAS-GFP*; *actP* > *CD2* > *LacZ*/+. For feeding experiments with bacteria or Dss, 2-3d old female adult flies were used for feeding experiments. Flies were cultured in an empty vial containing a piece of chromatography paper (Fisher) wet with 5% (wt/vol) sucrose (suc) solution as feeding medium (mock treatment) or with *P. e* or *Ecc15* or 5% Dss and 5% (wt/vol) sucrose for one or more days. For experiments involving tubGal80^ts^, the cross with right genotypes were set up and cultured at 18°C to restrict Gal4 activity. Two- to three-day-old F1 adult flies were shifted to 29°C for the indicated periods of time to inactivate Gal80^ts^ and allow Gal4 to activate UAS transgenes. For bacterial infection with knockdown of genes, flies bearing *Su(H)^ts^*>*GFP* or *Su(H)^ts^*>*EGFR-RNAi* (or *Ras-RNAi*)) were raised to adults at 18°C, and these adult females were transferred to 29°C for 6d and then fed with *P. e* for 24h. For the lineage tracing experiments with *P. e* infection, flies bearing *UAS-Flp*; *Su(H)^ts^UAS-GFP*; *actP*>*stop*>*LacZ* (or with *UAS-EGFR-RNAi*) were raised to adults at 18°C and then were transferred to 29°C for 6d and followed by feeding with suc (control) or *P. e* for 36h, and their midguts were immunostained with antibodies to GFP, β-gal and Dl (ISC marker) or Pros (EE marker). For two-color lineage tracing experiments, the cross with right genotypes were set up and cultured at 18 °C, and then 3-5-day-old flies expressing *UAS-Flp; Su(H)^ts^; FRT82BGFP/FRT82BRFP* with or without *UAS-gene* raised in 18°C was transferred to 29°C for the indicated time.

### Immunostaining

Female flies were used for gut immunostaining in all experiments. The entire intestine was dissected out and fixed in 1× PBS plus 8% EM-grade paraformaldehyde (Polysciences) for 1 h. Samples were washed and incubated with primary and secondary antibodies in a solution containing 1× PBS, 0.5% BSA, and 0.1% Triton X-100. The following primary antibodies were used: mouse anti-Delta (DSHB), 1:100; rabbit anti-LacZ (MP Biomedicals), 1:1,000; rabbit and mouse anti-PH3 (Millipore), 1:1,000; goat anti-GFP (Abcam), 1:1,000; Mouse anti-Pros (MR1A); Rabbit anti-Pdm1 (gift from X. Yang, Institute of Molecular and Cell Biology, Singapore).

Secondary antibodies conjugated to Alexa Fluor 546 donkey anti-mouse and anti-rabbit (Molecular Probes) and Alexa Fluor 633 donkey anti-mouse and anti-rabbit and 488 Donkey anti-goat (Jackson immunoresearch) were used at 1:400. Fluorescently labeled samples were counterstained with DAPI for visualization of DNA. Images were captured with a Zeiss LSM 800 confocal microscope and assembled in Adobe Photoshop.

### RT-qPCR

Total RNA was extracted from 10 female guts using RNeasy Plus Mini Kit (74134; Qiagen), and cDNA was synthesized using the iScript cDNA synthesis kit (Bio-Rad). RT-qPCR was performed using iQ SYBR Green System (Bio-Rad). Primer sequences used are: 5’-TCACACATTTAGTGGTGGAAG-3’ and 5’-TTGTGATGCTTGAATTGGTAA-3’ (for *vn*), 5’-CGTGTTTGGCAACAACAAGT-3’ and 5’-TGTGGCAATGCAGTTTAAGG-3’ (for *krn*), and 5’-CGCCCAAGAATGAAAGAGAG-3’ and 5-AGGTATGCTGCTGGTGGAAC-3’ (for *spi*). RpL11 was used as a normalization control. Relative quantification of mRNA levels was calculated using the comparative CT method.

### Genotypes for flies in each figure

Figure 1 (B-K’) *Su(H)-Gal4 UAS-GFP*/+

Figure 2 (B-C”) *Su(H)-Gal4 UAS-GFP*/+. (D) *Su(H)-Gal4 tub-Gal80^ts^ UAS-GFP/*+. (E) *Su(H)-Gal4 tub-Gal80^ts^ UAS-GFP/+; UAS-Ras-RNAi/UAS-Ras-RNAi*. (F) *Su(H)-Gal4 tub-Gal80^ts^ UAS-GFP/+; UAS-EGFR-RNAi/ UAS-EGFR-RNAi*.

Figure 3 (A-B’) *Su(H)-Gal4 tub-Gal80^ts^ UAS-GFP*/+. (C-E’) *Su(H)-Gal4 tub-Gal80^ts^ UAS-GFP/UAS-Ras^V12^*. (F-H’) *Su(H)-Gal4 tub-Gal80^ts^ UAS-GFP/UAS-EGFR^A887T^*. (I-K’) *Su(H)-Gal4 tub-Gal80^ts^ UAS-GFP/UAS-λTop*. (N-O’) *Su(H)-Gal4 tub-Gal80^ts^ UAS-GFP/+*. (P-R’) *Su(H)-Gal4 tub-Gal80^ts^ UAS-GFP/UAS-EGFR^A887T^*.

Figure 4 (B-C”’, E-E”) *UAS-Flp*; *Su(H)^ts^UAS-GFP; actP*>*stop*>*LacZ*. (F-F”’) *UAS-Flp*; *Su(H)^ts^UAS-GFP/UAS-Ras^V12^*; *actP*>*stop*>*LacZ*.

Figure 5 (A-A”, C-C”) *UAS-Flp*; *Su(H)^ts^UAS-GFP*; *actP*>*stop*>*LacZ*. (B-B”’) *UAS-Flp*; *Su(H)^ts^UAS-GFP/UAS-Ras^V12^*; *actP*>*stop*>*LacZ*.

Figure 6 (A-C) *UAS-Cnn-GFP*/+;; *Dl-Gal4 tub-Gal80^ts^*/+. (D-F) *UAS-Cnn-GFP*/+; *Su(H)-Gal4 tub-Gal80^ts^/UAS-EGFR^A887T^* (G-I) *UAS-Cnn-GFP/*+; *Su(H)-Gal4 tub-Gal80^ts^/UAS-Ras^V12^*. (L-L”) *UAS-Flp; Su(H) -Gal4 tub-Gal80^ts^; FRT82BGFP/FRT82BRFP*. (M-M”, O-O”) *UAS-Flp; Su(H) - Gal4 tub-Gal80^ts^/UAS-EGFR^A887T^; FRT82BGFP/FRT82BRFP*.

### Genotypes for flies in each supplementary figure

Figure 1—figure supplement 1. *Su(H)-Gal4 tub-Gal80^ts^ UAS-GFP/*+.

Figure 3—figure supplement 1. *Su(H)-Gal4 tub-Gal80^ts^ UAS-GFP/*+; *UAS-InR^ACT^/*+.

Figure 6—figure supplement 1. *UAS-Flp; Su(H)^ts^/UAS-EGFR^A887T^; FRT82BGFP/FRT82BRFP*.

## Acknowledgements

We thank Bloomington *Drosophila* Stock Center (NIH P400D018537), VDRC and Developmental Studies Hybridoma Bank for providing fly lines and antibodies, and thank Professors Yiping Chen, SM Jazwinski, Vivian Fonseca, Hong Liu and Jun-yuan Ji at Tulane University for discussion. This work was supported by the National Institutes of Health (grant P20GM103629 to A.T; grant GM072562, CA224381 and CA227789 to W.M. D.) and Carol Lavin Bernick Faculty Grant at Tulane University (A.T.).

---

**Figure 1—figure supplement 1.**
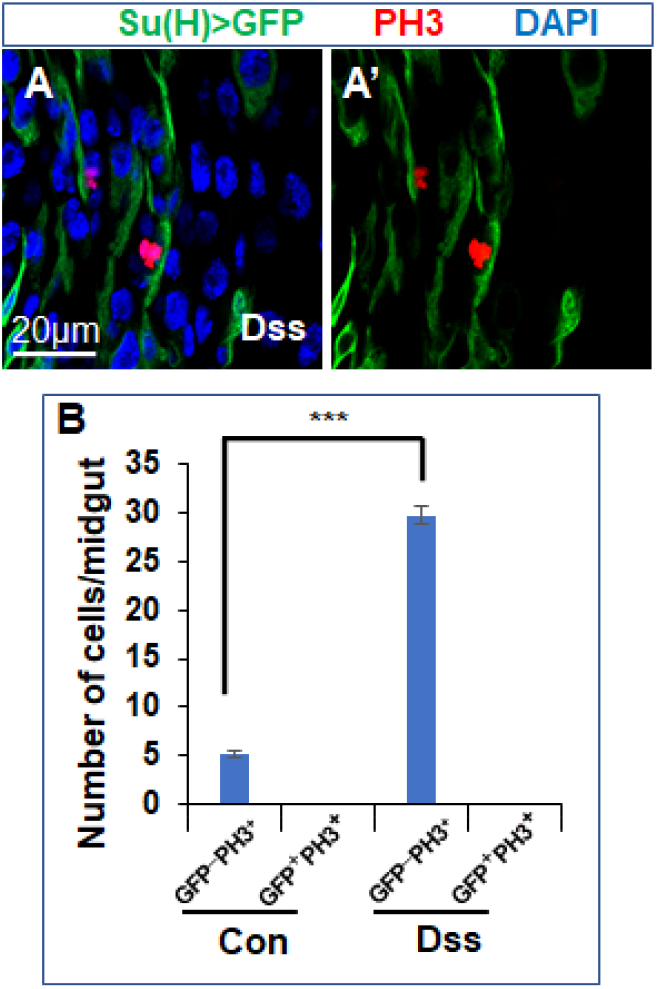
Feeding with Dss promotes ISC proliferation. (A-A’) The *Drosophila* midguts expressing *Su(H)^ts^*>*GFP* fed with Dss were immunostained for GFP (green), PH3 (red) and DAPI (blue). (B) Quantification of PH3^+^ in ISCs (GFP^−^) and EBs (GFP^+^). n = 8 guts for each genotype. Three independent experiments were performed, and error bars ± SEM. ***, P < 0.001 (Student’s t-test).

**Figure 3—figure supplement 1.**
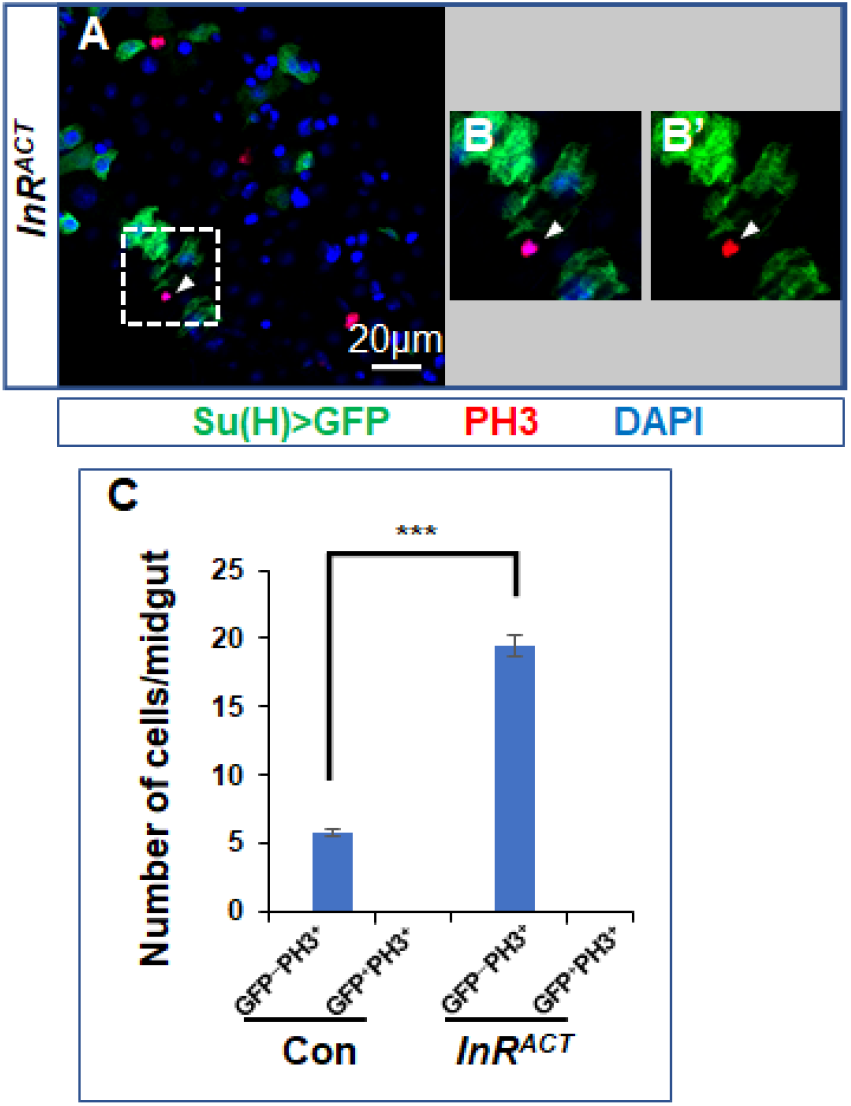
Activation of Insulin signaling with *InRA^CT^* in EBs non-cell-autonomously promotes ISC proliferation. (A) The *Drosophila* midguts expressing *Su(H)^ts^*>*GFP* with *InR^ACT^* for 4 days at 29°C were immunostained for GFP (green), PH3 (red) and DAPI (blue). (B-B’) Magnification of selecting areas containing PH3^−^GFP^−^ cells from A. (C) Quantification of PH3^+^ in ISCs (GFP^−^) and EBs (GFP^+^). n = 12 guts for each genotype. Three independent experiments were performed, and error bars ± SEM. ***, P < 0.001 (Student’s t-test).

**Figure 6—figure supplement 1.**
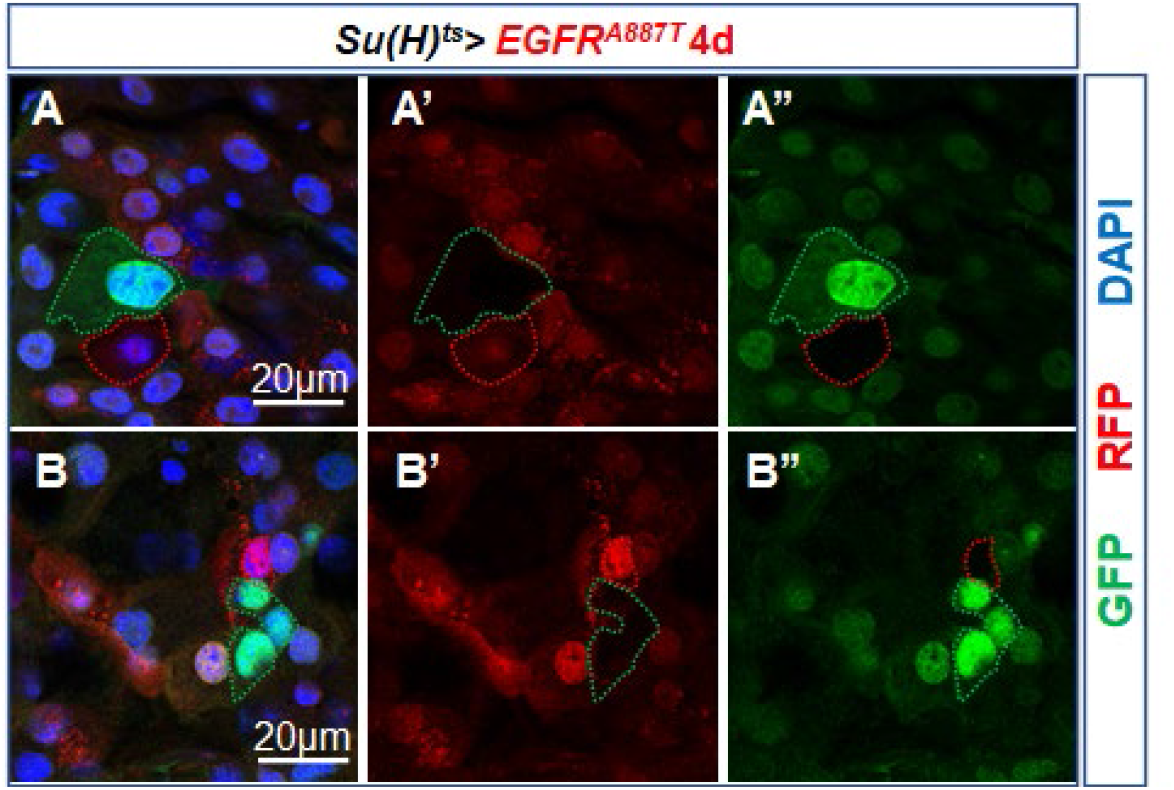
Activation of EGFR-Ras signaling in EBs induces EB dedifferentiation to generate one ISCs or non-ISCs. Representative twin clones from midguts shows symmetric division in EBs to produce two non-ISCs (A-A”) or asymmetric division in EB to produce one ISC with multiple cells (green) and one non-ISC (red) (B-B”).

